# Elucidating Interactome Dynamics of the A2A Adenosine Receptor

**DOI:** 10.1101/2024.07.08.601789

**Authors:** Wonseok Lee, Ahrum Son, Woojin Kim, Jongham Park, Ja-Young Cho, Ju-Won Kim, Young-Ok Kim, Hee Jeong Kong, Hyunsoo Kim

**Affiliations:** Department of Bio-AI convergence, Chungnam National University, 99 Daehak-ro, Yuseong-gu, Daejeon 34134, Republic of Korea; Department of Molecular Medicine, Scripps Research, La Jolla, CA 92037, United States; Biotechnology Research Division, National Institute of Fisheries Science, Busan 46083, Republic of Korea; Department of Convergent Bioscience and Informatics, Chungnam National University, 99 Daehak-ro, Yuseong-gu, Daejeon 34134, Republic of Korea; Protein AI Design Institute, Chungnam National University, 99 Daehak-ro, Yuseong-gu, Daejeon 34134, Republic of Korea; SCICS, Sciences for Panomics, 99 Daehak-ro, Yuseong-gu, Daejeon 34134, Republic of Korea

## Abstract

Polydeoxyribonucleotide (PDRN) is a tissue regeneration substance that resembles human DNA and is present in human cells, mullets, salmons, and flatfish. It stimulates physiological regeneration and metabolic activity. The regenerative and metabolic effects of PDRN are attributed to the activation of Adenosine A2A receptors (ADORA2A), which increases the production of angiogenesis factors and growth factors. Activation of ADORA2A leads to the activation of ADORA2A-interacting proteins with similar functions. To investigate the changes and dynamics of proteins in the presence of PDRN, we conducted selected reaction monitoring-mass spectrometry (SRM-MS) to quantify 491 proteins, 3,852 peptides. Through peptide-level analysis, we founded 374 proteins and 1,193 peptides demonstrating both up-regulation and down-regulation in expression. We conducted gene ontology (GO) analysis and physical network analysis and discovered a novel network of proteins related to neuronal differentiation among the ADORA2A-interacting protein. Based on network analysis, we found indirect interactions with MAPK1 and MP2K1, which are known to influence neuronal cell differentiation and suggest the formation of a network involving MAPK signaling transduction. Finally, using AlphaFold multimer, we were able to predict interaction sites among ADORA2A-interacting proteins in the network associated with neuronal cell differentiation. Specifically, we predicted five interaction sites between ADORA2A and NTRK1 (which interacts with ADORA2A), forming the edge. Thus, we provided indicators for further research using ADORA2A present in robust network and highlighted the potential of PDRN to impact neuronal differentiation.

## Introduction

Polydeoxyribonucleotide (PDRN) is a mixture of deoxyribonucleotides with molecular weights ranging from 50 to 1,500 KDa. It is obtained from human placentas and the semen of salmonid fish such as *Oncorhynchus mykiss* (rainbow trout) or *Oncorhynchus keta* (chum salmon)^(1)^. Although PDRN can be extracted from human placentas, drugs or beauty products containing PDRN are more commonly derived from the sperm of *Oncorhynchus mykiss* and *Oncorhynchus keta*^(3)^. PDRN serve as a DNA-derived drug providing purine and pyrimidine deoxynucleosides/deoxyribonucleotides and bases in both experimental and clinical studies^(2, 3)^. PDRN promotes vascular endothelial growth factor (VEGF) expression, angiogenesis, wound healing, osteoblast proliferation, anti-ischemic effect, and anti-inflammatory effects^(4–4)^. Efforts to understand the effective impact of PDRN are actively investigated in both *in vivo* and *in vitro* studies, focusing on optimal dosages and molecular sizes to maximize its efficacy^(9)^.

Activation of Adenosine A2A receptors (ADORA2A) is essential in the mechanisms that promote the self-regeneration of damaged cells and tissues involving PDRN^(1, 10)^. ADORA2A belongs to the superfamily of G protein-coupled receptors (GPCRs) with seven transmembrane domains, and its expression is observed in a wide variety of cells, including neurons, immune system cells, and muscle cells^(11)^. Activation of ADORA2A by adenosine leads to the activation of the protein kinase A (PKA)-mediated signal transduction pathway, which contributes to various physiological and pathological functions^(12)^. ADORA1 and ADORA2A receptors are G protein-coupled with activating or inactivating effects on adenylyl cyclase, which in turn affects cAMP, protein kinase A (PKA), subsequently regulating cellular functions^(13)^. Additionally, activated PKA influences downstream signaling pathways, including Ca^2+^ influx via the N-methyl-D-aspartate (NMDA) receptor, cyclic AMP (cAMP) generation, PKA activation, mitogen-activated protein kinase (MAPK) and cAMP-response element-binding protein (CREB) phosphorylation, and the subsequent transcription of plasticity-associated genes^(14)^. Its pathways serve as a crucial regulator in numerous physiological processes spanning neurotransmission, cardiac function, immune response modulation, and smooth muscle contraction regulation^(15)^. By activating ADORA2A, PDRN regulates the secretion of growth factors, promotes blood vessel expansion through VEGF, and exerts an anti-inflammatory effect^(16)^.

Various physiological and pathological processes occurring within cells are orchestrated through interactions among components such as proteins. Elements existing within networks with similar functions are referred to as the ‘interactome’^(17, 18)^. The interactome encompasses the entirety of molecular interactions within a cell, with a particular emphasis on protein-protein associations. These interactions define discursive networks that regulate various cellular processes, serving as critical components in understanding the complexity of biological systems^(19, 20)^. Interactions among these proteins occur through physical binding, elucidating which aids in a deeper understanding of the interactome^(21)^. Furthermore, unraveling these interactions aids in a more comprehensive understanding of the discursive signaling cascades mediated by these molecular associations. Understanding protein function relies on its physical interaction with partner proteins at specific binding sites, further enhancing our comprehension of cellular processes.

To date, the interaction between ADORA2A and its downstream partners following PDRN activation remains unexplored. In this study, we employed selected reaction monitoring-mass spectrometry (SRM-MS) to analyze the adenosine A2A receptor interactome in the presence of PDRNs isolated from *Paralichthys olivaceous* and *Acipenser gueldenstaedtii.* This approach allowed us to investigate quantitative changes in the ADORA2A interactome upon PDRN treatment. Furthermore, eludicating protein structure and predicting protein-protein interactions enable the identification of sites where these interactions occur. Finally, we provided additional indicators for research on neuronal differentiation by predicting new networks and interaction sites, thereby contributing to further studies in this field.

## Materials and Methods

### Sample Preparation

Genomic DNA was extracted using the TNES-urea buffer method^(22)^. Genomic DNA from the semen of *P. olivaceus* or *A. gueldenstaedtii* was incubated overnight at 55°C in lysis buffer (20 mM Tris-HCl (pH8.0), 20 mM EDTA, 200 mM NaCl, 4% SDS), 80 mM dithiothreitol (DTT), and proteinase K. Total genomic DNA (100 ng/μL) was sonicated with 30’’ ON/OFF pulses using an ultrasonic homogenizer ULH700S (Ulsso Hitech Co.). During sonication, the DNA sample was stored on ice. Samples sonicated for 18, 36 or 45 minutes were evenly mixed to produce DNA fragments below 1000 bp. The DNA solution was concentrated by ethanol precipitation to a final concentration of 10 μg/μL.

### Cell Culture and Treatment of PDRN

Human umbilical vein endothelial cells (HUVECs) were purchased from ATCC. HUVECs were plated onto 0.2% gelatin-coated plates and grown in sterile endothelial growth medium (EGM-2 BulletKit, Lonza) containing endothelial basal medium (EBM-2), 20% fetal bovine serum (FBS) (Gibco), 1% penicillin-streptomycin (Gibco), 3 ng/mL bFGF (Millipore), 5 U/mL heparin (Sigma). Cells were maintained at 37°C in a 5% CO_2_/95% air humidified atmosphere and subcultured when they were 70-80% confluent. The medium was changed every 48 hours. HUVECs at passage 3 were suspended in M199 culture medium (Gibco BRL, NY, USA) containing 5% FBS and plated onto a 100 mm dish coated with 0.2% gelatin. Cells were treated with PDRN to a final concentration of 100 μg/mL for 6 hours.

### Protein Extraction and Digestion

Protein samples were extracted from HUVECs and sonicated using 50 μL of 20 mM ammonium bicarbonate (ABC) with a Bioruptor set to ultra-high intensity for 30 minutes. The resulting supernatants were transferred to new tubes for BCA assay and subsequent digestion. Each tube, containing 100 μg of protein, underwent reduction of disulfide bonds in 2% sodium deoxycholate (SDC) and 10 mM tris-(2-carboxyethyl)-phosphine hydrochloride (TCEP) at 60℃ for 60 minutes. Cysteines were then alkylated using 20 mM 2-iodoacetamide (IAA) at 25℃ for 30 minutes in darkness. After diluting ABC to 20 mM with distilled water, proteins were digested with trypsin (Sequencing Grade Modified Trypsin, 20 μg, Promega) at an enzyme-to-protein ratio of 1:50 in 20 mM ABC, followed by overnight incubation at 37℃. The digestion reaction was terminated by adding 1% formic acid. The digested samples were combined with 5 μL of 6 x 5 LC-MS/MS Peptide Reference mix (Promega) and stored at-20℃ until analysis.

### Selected Reaction Monitoring-Mass Spectrometry (SRM-MS) Analysis

LC-MS/MS experiments were conducted using a 1290 Infinity Ⅱ Bio 2D-LC System (Agilent Technology), coupled with a 6495C Triple Quadrupole mass spectrometer (Agilent Technology) equipped with a Jet-stream electrospray source. Separation was achieved on ZORBAX Eclipse Plus C18 column (ID: 959759-902, length: 150 mm, pore size: 95 Å, particle size: 1.8 μm, Aglient Technology) using Mobile Phase A (0.1% formic acid in water) and Mobile Phase B (0.1% formic acid in acetonitrile). Samples (10 μg) were injected onto the column at a flow rate of 0.4 mL/min. SRM-MS data were processed using Skyline (MacCoss Lab), with additional analyses conducted in Excel (Microsoft Office 16).

### Statistical Analysis

The normalized intensity values of SRM-MS were calculated by normalizing to internal standard (IS) intensities for each method. *P*-values were derived from the Mann Whitney U test, comparing the intensities of transition groups between two different conditions using IBM SPSS statistics (version 27). Fold-changes (FC) were calculated by dividing the intensity of one group by that of the other using Excel (Microsoft). Proteins with transitions showing an absolute fold change greater than 2 and a *P*-value less than 0.05 were identified as differentially expressed proteins.

### GO Analysis & Network Analysis

Pathways associated with ‘neuronal differentiation’ were identified among the differentially expressed proteins through Gene Ontology (GO) analysis. We employed the DAVID tool for statistical analysis, applying the Benjamini and Hochberg False Discovery Rate (FDR) for correction, with a significance threshold of *P*-value < 0.05. Analyses were conducted across the categories of ‘biological process’, ‘cellular component’, and ‘molecular function’.

### Predicted Protein Structure & Interaction Site

To elucidate the structures of proteins implicated in neuronal differentiation, we employed structure prediction using AlphaFold Multimer (Version: 1.5.5) with the following setting: no template, amber, MSA mode set to mmseqs2_uniref_env, pair mode set to unpaired_paired, 20 cycles of recycling, and 5 cycles of relaxation. Sequence data were retrieved from Uniprot. Predictions of interactions among proximate proteins, identified through network analysis, were conducted using LocalFold (AlphaFold_batch.ipynb). The resulting predicted structures were visualized using PyMol (version 2.5.3). Interaction sites were predicted based on atoms within a 4 Å radius, forming hydrogen bonds or salt bridges between adjacent amino acids.

## Results

### Workflow of SRM-MS in ADORA2A Interactome Analysis

To investigate the alterations in protein composition of the ADORA2A interactome, we used a SRM-MS workflow to profile the dynamics and expression patterns of proteins interacting with ADORA2A in response to each PDRN from two different species (Flatfish and Salmon) and their different reproductive tissues (sperm and testis). Initially, we selected 513 proteins, 6,503 peptides known to interact with ADORA2A from an extensive literature review and public interactome databases, including STRING, HuRI, BioGRID, IntAct, MINT, HINT, Mentha, GPS-Prot, Wiki-Pi, and PIPs. Only peptides with detectable signals and unique mass values, identifiable using the SRM-MS analysis equipment, were selected, resulting in the identification of 491 proteins and 3,852 peptides. HUVECs were separately incubated for 6 hours in each of specific PDRN extracted from flatfish testis, sperm, salmon sperm (Fig. S1). The intensities obtained from the SRM-MS analysis were normalized using an Internal Standard (IS), and any missing values were imputed using the k-nearest neighborhood (kNN) algorithm. Fold-change (FC) and *P*-values were calculated based on the normalized values. 374 proteins demonstrating differential expression, as indicated by associated FC and *P*-value thresholds (|FC| > 2, *P*-value < 0.05), were further subjected to Gene Ontology (GO) analysis, network analysis, and structure prediction. These analytical steps were instrumental in providing a comprehensive and systematic understanding of the protein interactions (Fig. 1).

**Fig. 1.**
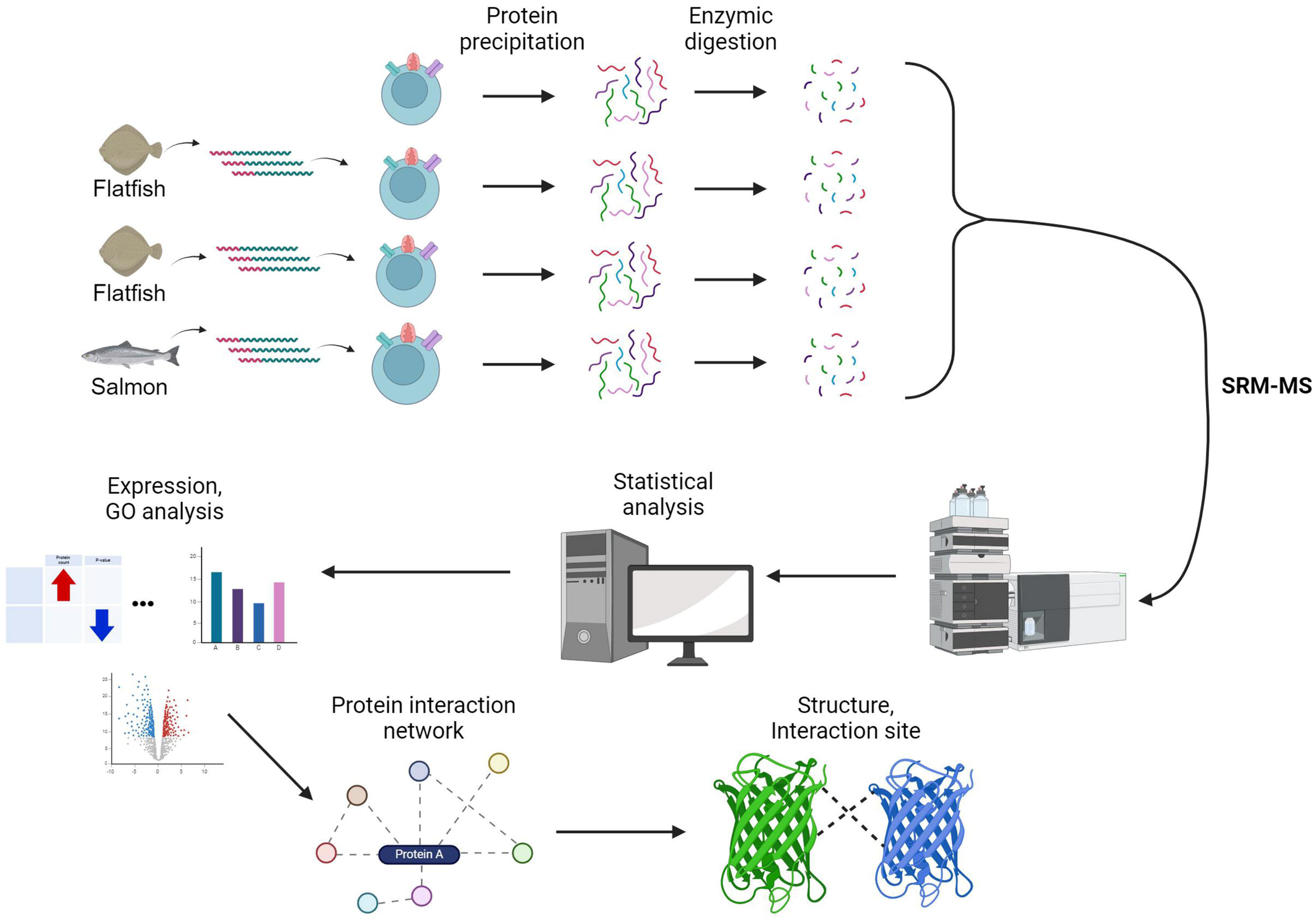
Workflow of Target Detection Interacting with ADORA2A in PDRN Environments. The initial identification of potential ADORA2A interactome targets was performed through an extensive literature review and public interactome databases (STRING, HuRI, BioGRID, IntAct, MINT, HINT, Mentha, GPS-Prot, Wiki-Pi, and PIPs), resulting in a preliminary list of 513 proteins and 6,503 peptides. Subsequently, unique protein markers were cataloged (502 proteins, 5,326 peptides). Using SRM-MS analysis, each sample was meticulously analyzed to detect the top two transitions in our experimental setup (491 proteins, 3,852 peptides, 7,704 transitions). The control group consisted of mock samples, while the experimental group included HUVEC cells incubated with salmon sperm, flatfish sperm, and flatfish testes for 6 hours. The experimental groups were designated as ‘Flatfish testis’ (HUVEC cells + flatfish testes), ‘Flatfish sperm’ (HUVEC cells + flatfish sperm), ‘Salmon sperm’ (HUVEC cells + salmon sperm), and the ‘Control group’ (HUVEC cells without PDRN).

### Comparative Analysis of the ADORA2A Interactome Across Different PDRN Species

We sought to investigate the expression patterns of the proteins that interact with ADORA2A depending on PDRN in different environments. To examine the changes in expression of ADORA2A-interacting proteins by PDRN in flatfish and salmon, and in their testis and sperm, we compared the number of proteins and peptides exhibiting an absolute fold change (|FC|) greater than 2 and *P*-values less than 0.05. The Mann-Whitney U test was employed for *P*-value determination (Fig. 2).

**Fig. 2.**
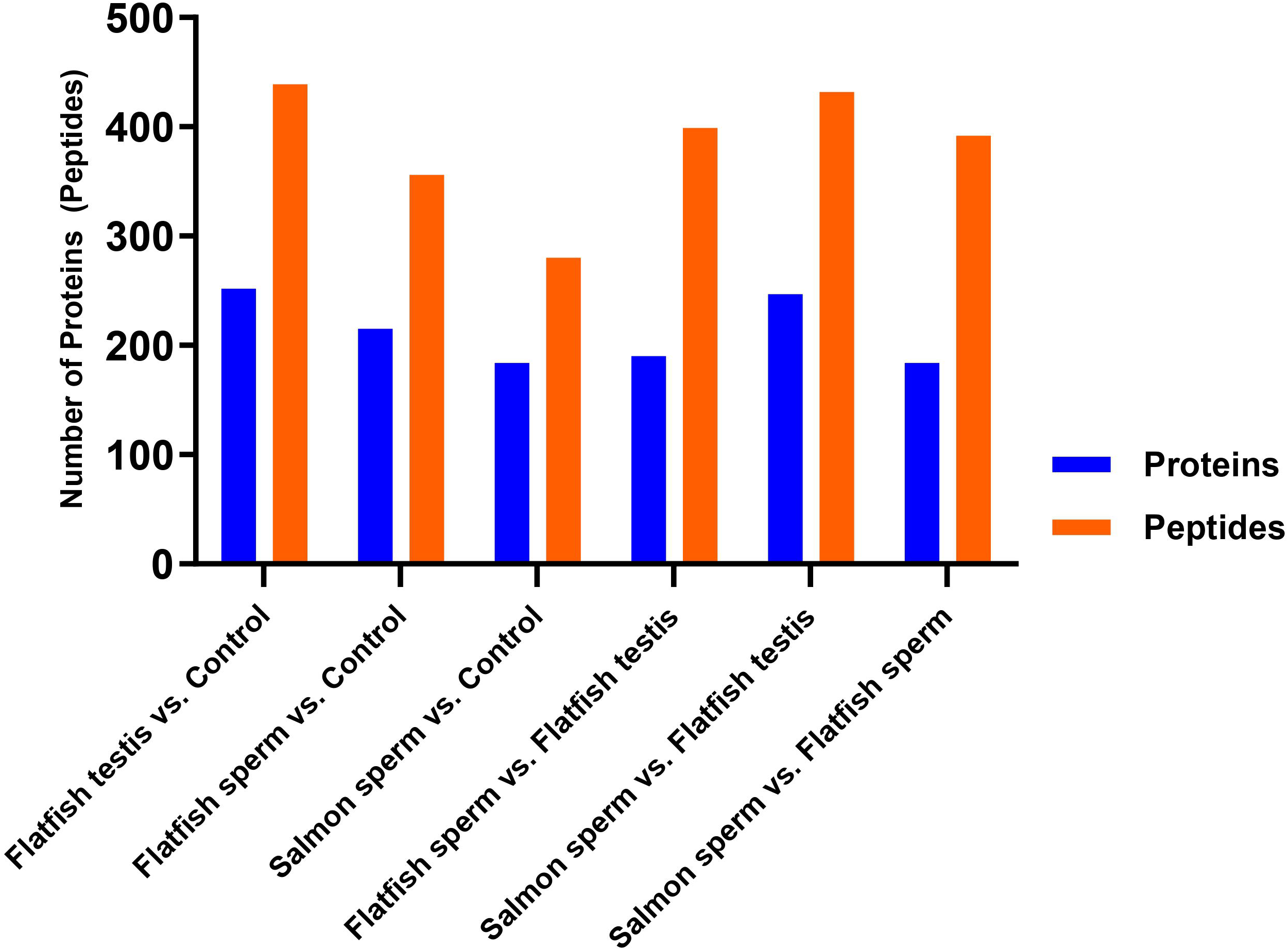
Comparative Analysis of the ADORA2A Interactome Across Different Groups Using Protein and Peptide Levels. This bar plot illustrates the number of differentially expressed proteins and peptides between two groups based on the criteria of |FC| > 2 and a *P*-value < 0.05. Differentially expressed proteins are depicted in blue, while peptides are shown in orange. The fold change (FC) was calculated by dividing the expression levels in the group before ‘vs’ by the group after ‘vs’.

The most differential proteins were exhibited in the sample with the testis of the flatfish, where 252 proteins and 439 peptides were significantly differentially expressed compared to the control group. When treated with salmon sperm, the fewest proteins–184 proteins and 280 peptides–were differentially expressed compared to the control group. These findings revealed that various types of PDRN influence the expression levels of the ADORA2A interactome, and this influence varies depending on the type of PDRN.

Next, we quantified the change in the expression of the differential proteins across groups. Normalized intensities with FC greater than 2 and *P*-value less than 0.05 were classified as ‘Up-regulation’, whereas those with FC less than 0.5 and *P*-value less than 0.05 were categorized as ‘Down-regulation’. There were 192 proteins and 268 peptides showing the highest number of up-regulation targets in ‘Salmon sperms vs. Flatfish sperms’ (Fig. S2A) and 138 proteins and 189 peptides showing the highest number of down-regulation targets in ‘Flatfish sperm vs. Control group’ (log_2_(FC) > 1 or <-1, *P*-value < 0.05) (Fig. S2B). In the ‘Flatfish testis vs. Control group’, 304 proteins and 401 peptides exhibited differentially expressed targets, including MAPK1 (Mitogen-activated protein kinase 1) and MAP2K1 (Dual specificity mitogen-activated protein kinase kinase 1), which are associated with signal transduction (Fig. S2C). The comparison between groups revealed variability in the number of intensities showing up-regulation and down-regulation. It suggests that the expression levels of ADORA2A-interacting proteins vary depending on the PDRN from the species and reproductive tissues from which they originated. (Fig. S3A-C).

### Network of Differentially Expressed ADORA2A Interactome Associated with Neuronal Cell Differentiation

To elucidate the functions in which the differentially expressed ADORA2A-interacting proteins are involved, we conducted a Gene Ontology (GO) analysis for each group (Table S1-6). The GO categories ‘Biological Processes’, ‘Cellular Components’, and ‘Molecular Functions’ were analyzed to identify enrichment patterns associated with the ADORA2A interactome proteins in response to PDRN. This analysis was facilitated by the DAVID tool focusing on GO terms with a *P*-value less than 0.05 to pinpoint the most pertinent differentially expressed proteins. The differential proteins in the ‘Flatfish testis vs. Control group’ comparison were significantly involved in processes such as signal transduction and protein phosphorylation (Table S1). Specifically, in the ‘Biological Processes’ category, proteins were predominantly linked to the signal transduction pathways, whereas in the ‘Cellular Components’ category, they were primarily associated with the plasma membrane. In the ‘Molecular Function’ category, the primary associations were with protein binding and ATP binding. These GO terms were associated with processes such as glial cell proliferation and growth, resonating with patterns observed in neuronal differentiation^(23)^.

To uncover the intricate interactions within the differentially expressed ADORA2A interactome (FC > 2, *P*-value < 0.05), we generated protein-protein interaction (PPI) networks for each group comparison using the STRING database (https://string-db.org/) to unravel the regulatory patterns among the comparison groups. In the network of proteins, we analyzed a physical subnetwork to identify the physical interactions among differentially expressed proteins related to neuronal cell differentiation (Fig. S4A-E). In the ‘Flatfish testis vs. Control group’ comparison, we identified 178 physical interactions among 137 unique proteins related to ADORA2A (confidence score > 0.7) (Fig. 3A).

**Fig. 3.**
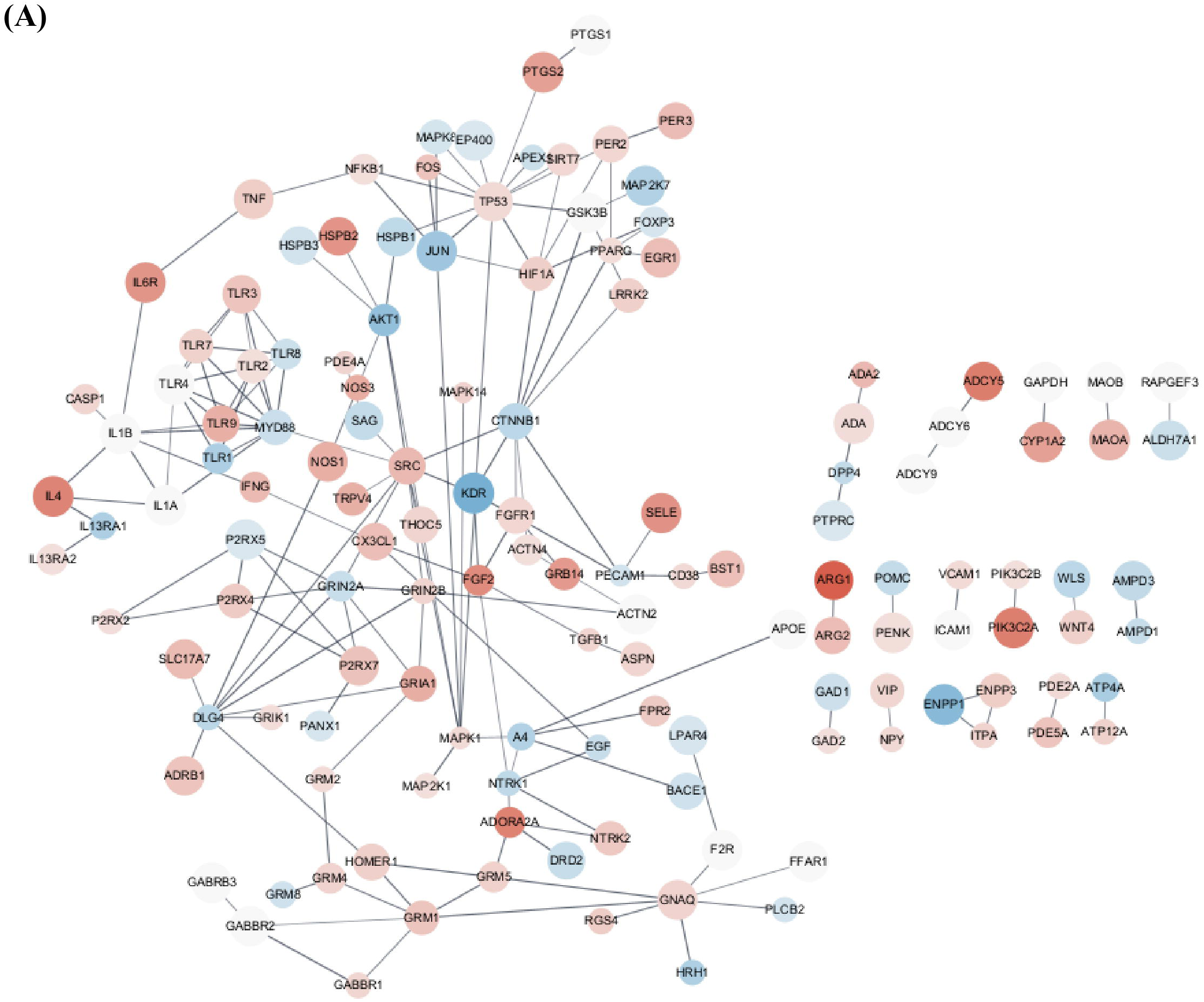
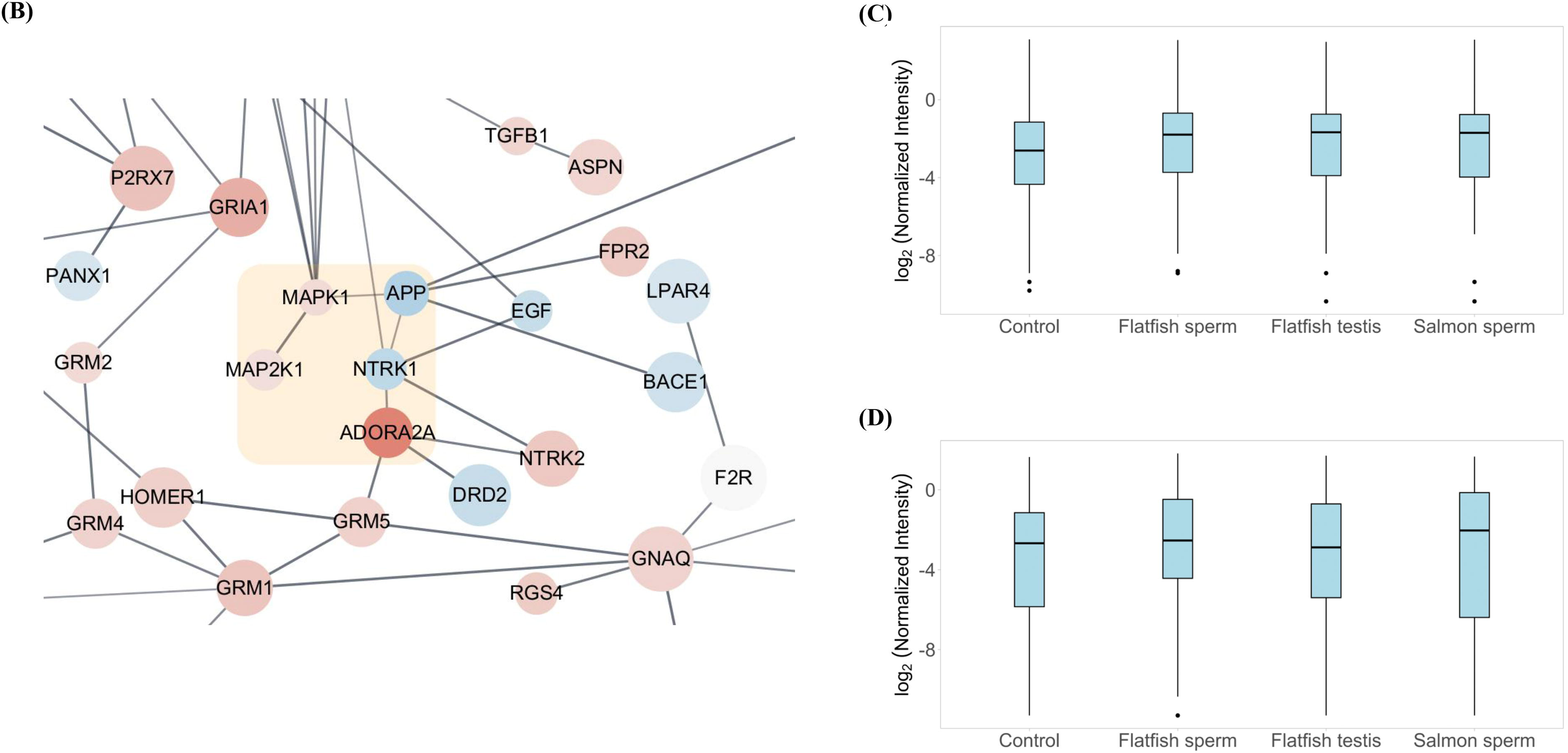
Participation of ADORA2A Interactome Proteins in Signal Transduction Associated with Neuronal Differentiation. (A) Network analysis depicting the interactions between ADORA2A and various proteins in the ‘Flatfish testis/Control group’, with red and blue colors indicating up-regulation and down-regulation, respectively. The size of each node reflects the *P*-value associated with that protein. (B) Network analysis highlighting the interactions between ADORA2A and proteins implicated in neuronal differentiation within the ‘Flatfish testis/Control group’ context. (C-D) Box plots representing the normalized intensities of MAPK1(up) and MAP2K1(down) across each group.

These interactions were visualized using Cytoscape software, displaying clusters of ADORA2A-interacting proteins and several key components involved in glial cell proliferation and kinase signaling pathways (Fig. 3B). We noted proteins such as NTRK1 (neurotrophic receptor tyrosine kinase 1), APP (Amyloid-beta precursor protein), MAPK1, and MAP2K1. These proteins play pivotal roles in signal transduction pathways and glial cell differentiation^(24–24)^. Despite variability in the expression profiles across species, the data suggested that PDRN might exert distinct influences on the MAP kinase pathway, thereby affecting ADORA2A interactions and neuronal cell differentiation (Fig. 3C, 3D).

### Predicted Interaction Sites in ADORA2A Interactome Associated with Neuronal Differentiation

In our network analysis for the comparison of ‘Flatfish testis vs. Control group’, we investigated the structures of ADORA2A-interacting proteins by predicting the interaction of the proteins and identifying their interaction sites. We selected proteins for their roles in regulating glial and neuronal cell differentiation. The comparison of ‘Flatfish testis vs. Control group’ was chosen because its network has physical interactions of proteins associated with neuronal differentiation. Using protein sequence information from UniProt, we predicted the structures of interacting proteins using AlphaFold-multimer, allowing us to dynamically represent protein interactions within the ‘Flatfish testis vs. Control group’ network (Fig. 4A-E). Using these structures, we predicted interaction sites between pairs of proteins based on their proximity. For instance, interaction sites between ADORA2A and NTRK1 were identified, marked by five binding interaction sites between ADORA2A and NTRK1 (Fig. 4B-D). Based on the results, we measured distances between proteins implicated in neuronal differentiation, identifying several interaction regions (Table 1). Interaction sites were colored magenta between NTRK1 and APP (Fig. 4E), and between APP and MAPK1 (Fig. 4F), and between MAPK1 and MP2K1 (Fig. 4G), highlighting the intricate nature of these interactions. Additionally, we discovered the highest number of predicted interaction sites between NTRK1 and APP. These predicted interaction regions, derived from sequence information, provided insights into the potential roles of differentially expressed proteins in neural differentiation within the comparison of ‘Flatfish testis vs. Control group’.

**Fig. 4.**
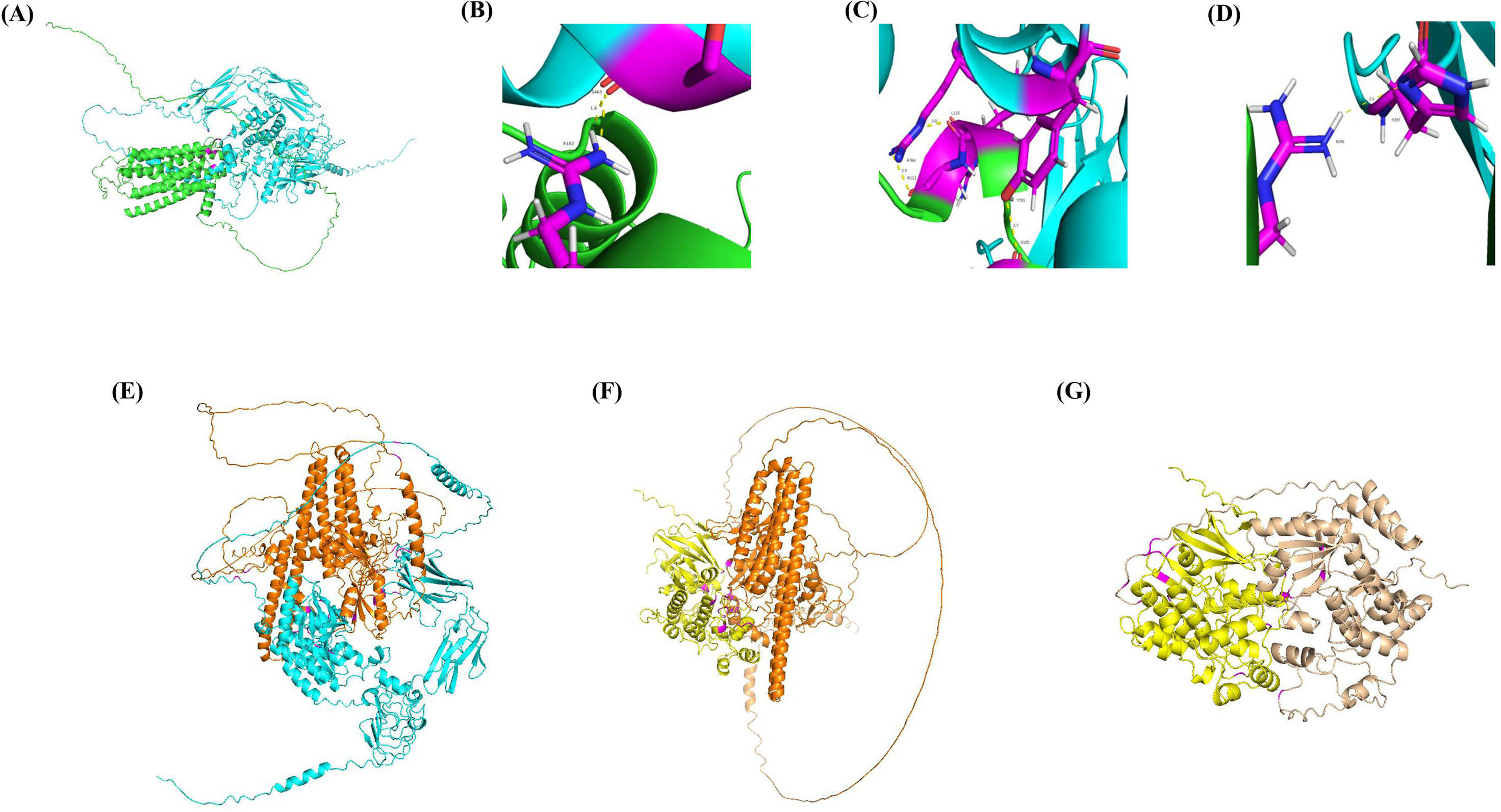
Predicted Structures and Interaction Sites of Network Proteins Associated with Neuronal Differentiation. (A) Composite structure of ADORA2A (green) and NTRK1 (cyan). (B-D) Highlighted interaction site (magenta) between ADORA2A (green) and NTRK1 (cyan) with stick model (ADORA2A: Arg206, Ala105, Arg111, Leu110, Arg102; NTRK1: His297, Tyr701, Arg766, Ser465). (E) Composite structure of NTRK1 (cyan) and APP (orange). (F) Composite structure of APP (orange) and MAPK1 (yellow). (G) Composite structure of MAPK1 (yellow) and MAP2K1 (apricot). Each interaction site is highlighted in magenta.

**Table 1.**
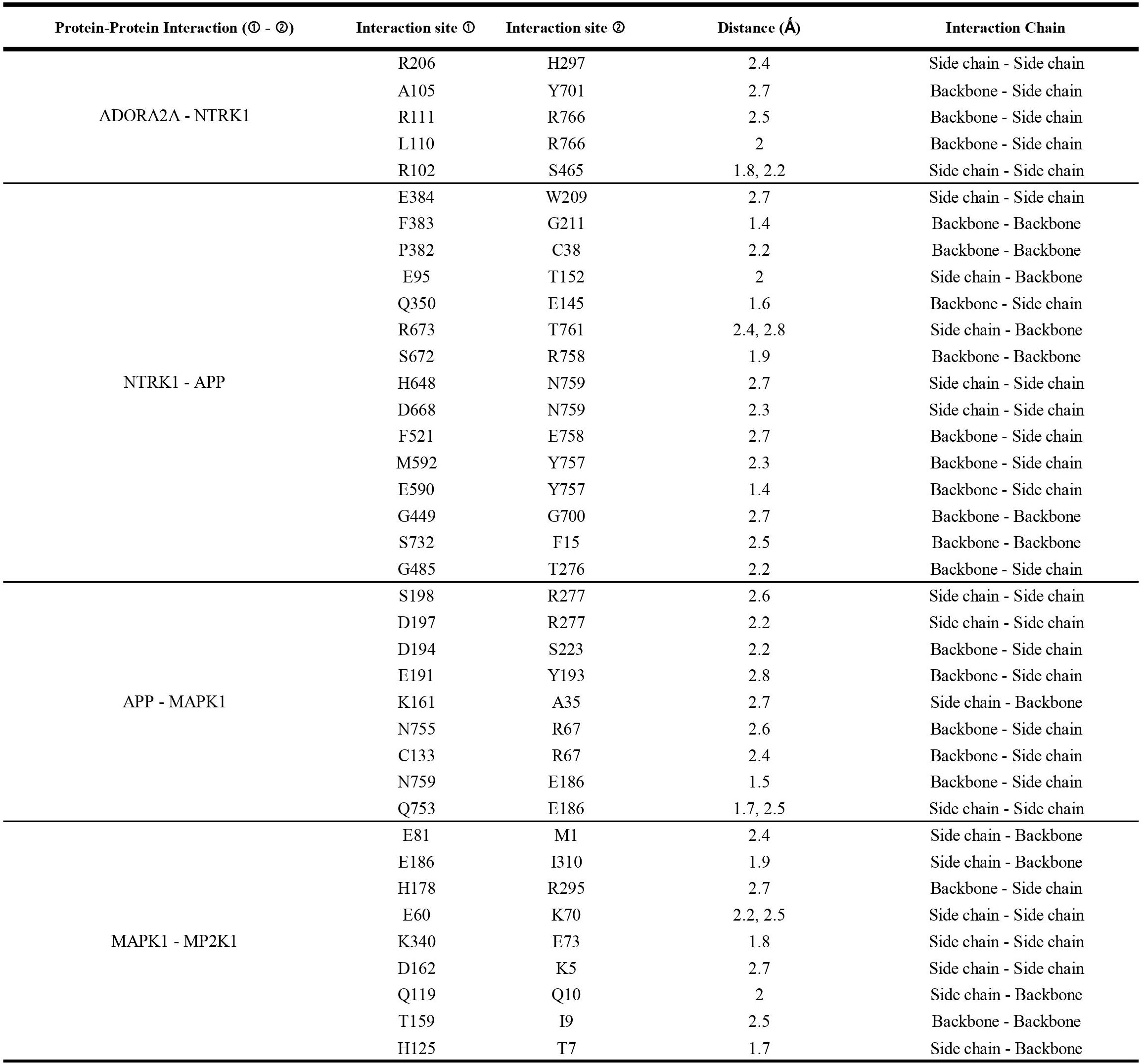
Predicted Interaction Site of Proteins Participating in Network Analysis of the ‘Flatfish testis/Control group’.

## Discussion

Our comprehensive workflow and detailed analyses offer a profound understanding of the ADORA2A interactome and its modulation in various PDRN environments. We used SRM-MS to identify ADORA2A-interacting proteins and gain insight into the biological functions of the proteins associated with neuronal cell differentiation. Our results revealed that activation of ADORA2A through PDRN can induce the regulation of ADORA2A-interacting proteins. The insights gleaned from GO analysis, PPI networks, and interaction site predictions contribute significantly to our knowledge of protein interactions and their implications in neuronal differentiation and related biological processes.

Further analysis of the protein-protein interaction network for ADORA2A-interacting proteins showed that in the ‘Flatfish testis vs. Control group’, proteins associated with ADORA2A were involved in signal transduction, such as MAPK signaling and neuronal differentiation. Proteins in the network are known to affect neurological diseases along with signal transduction and neural changes, which is consistent with previous research^(30–30)^.

Previous studies have demonstrated a negative correlation between the expression levels of ADORA2A and NTRK1 in NS (NeuroScreen) cells^(33)^. Our results exhibit a similar pattern, suggesting that the activation of ADORA2A through PDRN may influence NTRK1 expression and indicate a more accurate interaction within the physical subnetwork. Additionally, research has shown that in Alzheimer’s disease patients and advanced aged individuals, there is a mitochondrial accumulation of APP^(34)^. Conversely, in our activated network, APP was found to be downregulated, highlighting PDRN as a potential biomarker for neuro-related diseases, aligning with the observed downregulation of proteins within the activated network.

Likewise, MAPK signaling pathways have been implicated in the pathogenesis of a variety of diseases, including cancer and neurodegenerative diseases such as Alzheimer’s disease^(35)^. Moreover, MAPK1, the key regulator of this pathway, is known to be a major mediator in excessive cell growth, such as in several cancer^(36–36)^. Our results identified interactions between ADORA2A and MAPK1, suggesting that this interaction network could serve as an indicator for studying cell tumorigenesis. While previous studies have investigated MAPK1 and ADORA2A individually^(39)^, our research distinguishes itself by uncovering the integrated network involving MAPK1 through activated ADORA2A. This finding provides a novel perspective for cancer research and highlights potential biomarkers for tumor growth and cell differentiation.

Despite discovering a network of ADORA2A-interacting proteins related to neuronal differentiation, our study had several limitations. Firstly, the number of replicates per group was limited. Although we used six replicates per group in this study, the results did not meet the normality assumption, preventing the use of adjusted *P*-values. This limitation highlights the need for further studies to include a larger number of replicates to ensure more robust and statistically valid findings. Secondly, it was not possible to determine the differential expression of neurodifferentiation-related proteins across all groups. Our study only identified the network of ADORA2A and its interacting proteins in the comparison of the ‘Flatfish testis vs. Control group’. Additionally, although it is known that receptors like DRD2 (D (2) dopamine receptor) are inhibited when ADORA2A is activated^(40)^, we observed such changes only in some group comparisons. This further underscores the necessity for more comparative studies to validate these interactions across different conditions and groups. Finally, our study is limited by the fact that we only predicted the interaction sites. The predictions were made using AlphaFold, a model that predicts protein structures based on their sequences^(41)^. However, in our prediction of interaction sites between ADORA2A and proteins involved in MAPK signaling, no interaction sites were observed (Fig. S5). Therefore, advancements in modeling or experimental validation are essential to confirm the precise interaction sites.

In conclusion, our comprehensive workflow and detailed analyses provide a profound understanding of the ADORA2A interactome and its modulation in various PDRN environments. The insights obtained from GO analysis, PPI networks, and interaction site predictions significantly enhance our knowledge of protein interactions and their implications in neuronal differentiation and related biological processes.

## Supporting information

ADORA2A_supplementary_data

## Acknowledgements

This work was supported by National Institute of Fisheries Science, Ministry of Oceans and Fisheries (R2023019). This work was also supported by Institute of Information & communications Technology Planning & Evaluation (IITP) grant funded by the Korea government (MSIT) (No.RS-2022-00155857, Artificial Intelligence Convergence Innovation Human Resources Development program at Chungnam National University. The authors declare that there are no conflicts of interest related to this research.

## References

1. Squadrito, F., Bitto, A., Irrera, N., Pizzino, G., Pallio, G., Minutoli, L., and Altavilla, D. (2017) Pharmacological activity and clinical use of PDRN. Frontiers in pharmacology 8, 224

2. Altavilla, D., Bitto, A., Polito, F., Marini, H., Minutoli, L., Stefano, V. D., Irrera, N., Cattarini, G., and Squadrito, F. (2009) Polydeoxyribonucleotide (PDRN): a safe approach to induce therapeutic angiogenesis in peripheral artery occlusive disease and in diabetic foot ulcers. Cardiovascular & Hematological Agents in Medicinal Chemistry (Formerly Current Medicinal Chemistry-Cardiovascular & Hematological Agents) 7, 313–321

3. Lee, J. H., Han, J. W., Byun, J. H., Lee, W. M., Kim, M. H., and Wu, W. H. (2018) Comparison of wound healing effects between Oncorhynchus keta-derived polydeoxyribonucleotide (PDRN) and Oncorhynchus mykiss-derived PDRN. Archives of Craniofacial Surgery 19, 20

4. Galeano, M., Bitto, A., Altavilla, D., Minutoli, L., Polito, F., Calò, M., Lo Cascio, P., Stagno d’Alcontres, F., and Squadrito, F. (2008) Polydeoxyribonucleotide stimulates angiogenesis and wound healing in the genetically diabetic mouse. Wound Repair and Regeneration 16, 208–217

5. Baek, A., Kim, Y., Lee, J. W., Lee, S. C., and Cho, S.-R. (2018) Effect of polydeoxyribonucleotide on angiogenesis and wound healing in an in vitro model of osteoarthritis. Cell transplantation 27, 1623–1633

6. Guizzardi, S., Galli, C., Govoni, P., Boratto, R., Cattarini, G., Martini, D., Belletti, S., and Scandroglio, R. (2003) Polydeoxyribonucleotide (PDRN) promotes human osteoblast proliferation: a new proposal for bone tissue repair. Life sciences 73, 1973–1983

7. Bitto, A., Polito, F., Altavilla, D., Minutoli, L., Migliorato, A., and Squadrito, F. (2008) Polydeoxyribonucleotide (PDRN) restores blood flow in an experimental model of peripheral artery occlusive disease. Journal of Vascular Surgery 48, 1292–1300

8. Han, J. H., Jung, J., Hwang, L., Ko, I. G., Nam, O. H., Kim, M. S., Lee, J. W., Choi, B. J., and Lee, D. W. (2018) Anti-inflammatory effect of polydeoxyribonucleotide on zoledronic acid-pretreated and lipopolysaccharide-stimulated RAW 264.7 cells. Experimental and therapeutic medicine 16, 400–405

9. Hwang, K. H., Kim, J. H., Park, E. Y., and Cha, S. K. (2018) An effective range of polydeoxyribonucleotides is critical for wound healing quality. Molecular Medicine Reports 18, 5166–5172

10. Thellung, S., Florio, T., Maragliano, A., Cattarini, G., and Schettini, G. (1999) Polydeoxyribonucleotides enhance the proliferation of human skin fibroblasts: involvement of A2 purinergic receptor subtypes. Life sciences 64, 1661–1674

11. Rahimi, M. R., Semenova, E. A., Larin, A. K., Kulemin, N. A., Generozov, E. V., Łubkowska, B., Ahmetov, I. I., and Golpasandi, H. (2023) The ADORA2A TT genotype is associated with anti-inflammatory effects of caffeine in response to resistance exercise and habitual coffee intake. Nutrients 15, 1634

12. Sheth, S., Brito, R., Mukherjea, D., Rybak, L. P., and Ramkumar, V. (2014) Adenosine receptors: expression, function and regulation. International journal of molecular sciences 15, 2024–2052

13. Fredholm, B. B., Abbracchio, M. P., Burnstock, G., Daly, J. W., Harden, K. T., Jacobson, K. A., Leff, P., and Williams, M. (1994) VI. Nomenclature and classification of purinoceptors. Pharmacological reviews 46, 143

14. Waltereit, R., and Weller, M. (2003) Signaling from cAMP/PKA to MAPK and synaptic plasticity. Molecular neurobiology 27, 99–106

15. Tang, Z., Ye, W., Chen, H., Kuang, X., Guo, J., Xiang, M., Peng, C., Chen, X., and Liu, H. (2019) Role of purines in regulation of metabolic reprogramming. Purinergic signalling 15, 423–438

16. Colangelo, M. T., Galli, C., and Guizzardi, S. (2020) The effects of polydeoxyribonucleotide on wound healing and tissue regeneration: a systematic review of the literature. Regenerative medicine 15, 1801–1821

17. Vidal, M., Cusick, M. E., and Barabási, A.-L. (2011) Interactome networks and human disease. Cell 144, 986–998

18. Cusick, M. E., Klitgord, N., Vidal, M., and Hill, D. E. (2005) Interactome: gateway into systems biology. Human molecular genetics 14, R171–R181

19. Lage, K. (2014) Protein–protein interactions and genetic diseases: the interactome. Biochimica et Biophysica Acta (BBA)-Molecular Basis of Disease 1842, 1971–1980

20. De Las Rivas, J., and Fontanillo, C. (2010) Protein–protein interactions essentials: key concepts to building and analyzing interactome networks. PLoS computational biology 6, e1000807

21. Vakser, I. A. (2014) Protein-protein docking: From interaction to interactome. Biophysical journal 107, 1785–1793

22. Asahida, T., Kobayashi, T., Saitoh, K., and Nakayama, I. (1996) Tissue preservation and total DNA extraction form fish stored at ambient temperature using buffers containing high concentration of urea. Fisheries science 62, 727–730

23. Magistri, M., Khoury, N., Mazza, E. M. C., Velmeshev, D., Lee, J. K., Bicciato, S., Tsoulfas, P., and Faghihi, M. A. (2016) A comparative transcriptomic analysis of astrocytes differentiation from human neural progenitor cells. European Journal of Neuroscience 44, 2858–2870

24. Wang, L., He, F., Zhong, Z., Lv, R., Xiao, S., and Liu, Z. (2015) Overexpression of NTRK1 promotes differentiation of neural stem cells into cholinergic neurons. BioMed Research International 2015

25. Pajtler, K. W., Mahlow, E., Odersky, A., Lindner, S., Stephan, H., Bendix, I., Eggert, A., Schramm, A., and Schulte, J. H. (2014) Neuroblastoma in dialog with its stroma: NTRK1 is a regulator of cellular cross-talk with Schwann cells. Oncotarget 5, 11180

26. Bergström, P., Agholme, L., Nazir, F. H., Satir, T. M., Toombs, J., Wellington, H., Strandberg, J., Bontell, T. O., Kvartsberg, H., and Holmström, M. (2016) Amyloid precursor protein expression and processing are differentially regulated during cortical neuron differentiation. Scientific reports 6, 29200

27. Coronel, R., Lachgar, M., Bernabeu-Zornoza, A., Palmer, C., Domínguez-Alvaro, M., Revilla, A., Ocaña, I., Fernández, A., Martínez-Serrano, A., and Cano, E. (2019) Neuronal and glial differentiation of human neural stem cells is regulated by amyloid precursor protein (APP) levels. Molecular neurobiology 56, 1248–1261

28. Sun, Y., Liu, W.-Z., Liu, T., Feng, X., Yang, N., and Zhou, H.-F. (2015) Signaling pathway of MAPK/ERK in cell proliferation, differentiation, migration, senescence and apoptosis. Journal of Receptors and Signal Transduction 35, 600–604

29. Kennedy, K. E., Kerlero de Rosbo, N., Uccelli, A., Cellerino, M., Ivaldi, F., Contini, P., De Palma, R., Harbo, H. F., Berge, T., and Bos, S. D. (2024) Multiscale networks in multiple sclerosis. PLOS Computational Biology 20, e1010980

30. Alberti, L., Carniti, C., Miranda, C., Roccato, E., and Pierotti, M. A. (2003) RET and NTRK1 proto-oncogenes in human diseases. Journal of cellular physiology 195, 168–186

31. Schettini, G., Govoni, S., Racchi, M., and Rodriguez, G. (2010) Phosphorylation of APP-CTF-AICD domains and interaction with adaptor proteins: signal transduction and/or transcriptional role– relevance for Alzheimer pathology. Journal of neurochemistry 115, 1299–1308

32. Deng, Y., Zhang, J., Sun, X., Ma, G., Luo, G., Miao, Z., and Song, L. (2020) miR-132 improves the cognitive function of rats with Alzheimer’s disease by inhibiting the MAPK1 signal pathway. Experimental and Therapeutic Medicine 20, 1–1

33. Pokharel, S., Lee, C. H., Gilyazova, N., and Ibeanu, G. C. (2018) Analysis of gene expression and neuronal phenotype in neuroscreen-1 (NS-1) cells. International journal of biomedical investigation 1

34. Pavlov, P. F., Petersen, C. H., Glaser, E., and Ankarcrona, M. (2009) Mitochondrial accumulation of APP and Aβ: significance for Alzheimer disease pathogenesis. Journal of cellular and molecular medicine 13, 4137–4145

35. Kim, E. K., and Choi, E.-J. (2010) Pathological roles of MAPK signaling pathways in human diseases. Biochimica et Biophysica Acta (BBA)-Molecular Basis of Disease 1802, 396–405

36. Saklatvala, J. (2004) The p38 MAP kinase pathway as a therapeutic target in inflammatory disease. Current opinion in pharmacology 4, 372–377

37. Li, X.-W., Tuergan, M., and Abulizi, G. (2015) Expression of MAPK1 in cervical cancer and effect of MAPK1 gene silencing on epithelial-mesenchymal transition, invasion and metastasis. Asian Pacific journal of tropical medicine 8, 937–943

38. Wang, Y., Guo, Z., Tian, Y., Cong, L., Zheng, Y., Wu, Z., Shan, G., Xia, Y., Zhu, Y., and Li, X. (2023) MAPK1 promotes the metastasis and invasion of gastric cancer as a bidirectional transcription factor. BMC cancer 23, 959

39. Hutton, S. R., Otis, J. M., Kim, E. M., Lamsal, Y., Stuber, G. D., and Snider, W. D. (2017) ERK/MAPK signaling is required for pathway-specific striatal motor functions. Journal of Neuroscience 37, 8102–8115

40. Childs, E., Hohoff, C., Deckert, J., Xu, K., Badner, J., and De Wit, H. (2008) Association between ADORA2A and DRD2 polymorphisms and caffeine-induced anxiety. Neuropsychopharmacology 33, 2791–2800

41. Evans, R., O’Neill, M., Pritzel, A., Antropova, N., Senior, A., Green, T., Žídek, A., Bates, R., Blackwell, S., and Yim, J. (2021) Protein complex prediction with AlphaFold-Multimer. biorxiv, 2021.2010. 2004.463034

