## Supplementary material for "Elucidating Interactome Dynamics of the A2A Adenosine Receptor": ADORA2A_supplementary_data: Supplementary_figure.pptx

### Slide 1
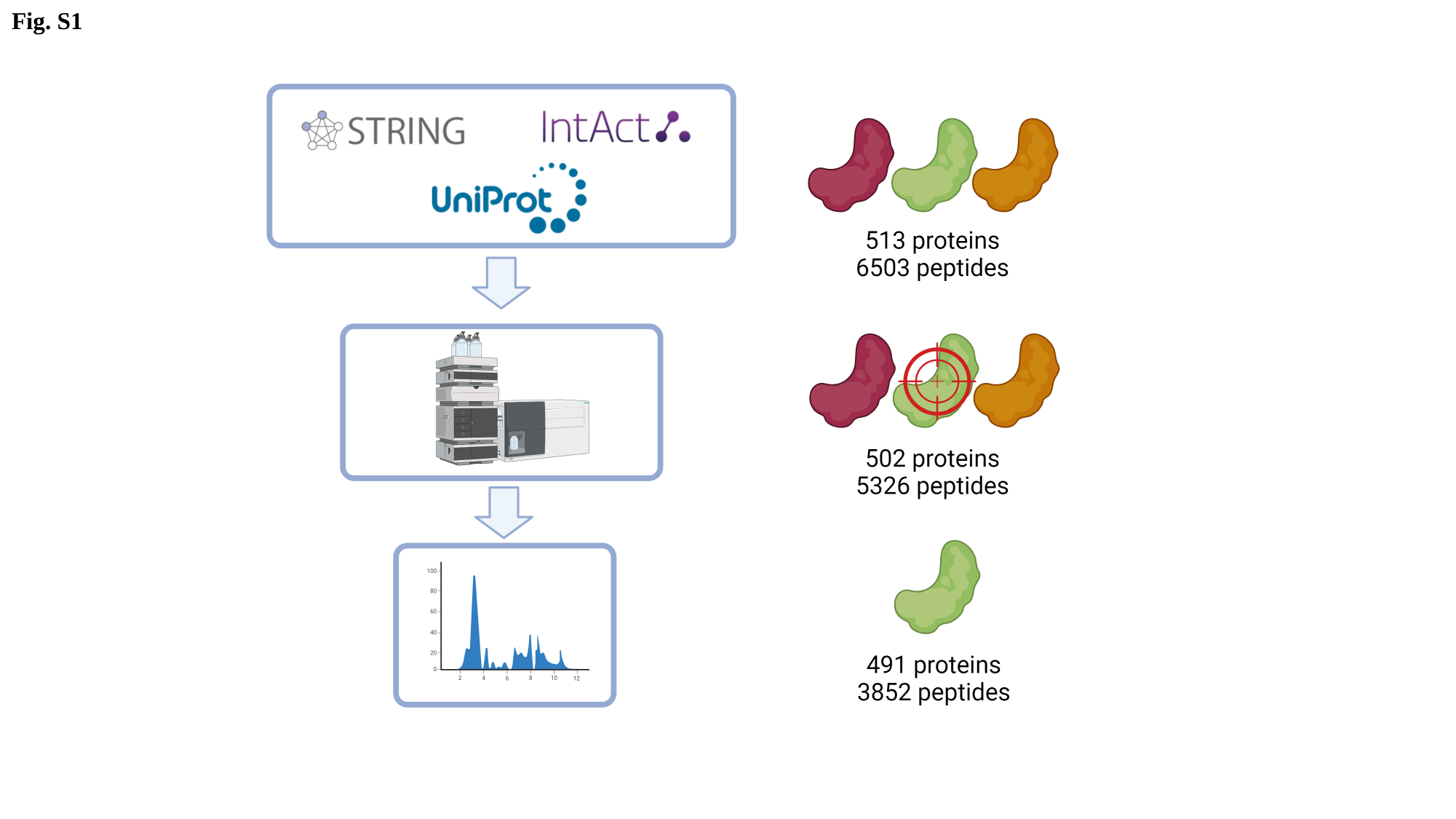

Fig. S1

### Slide 2
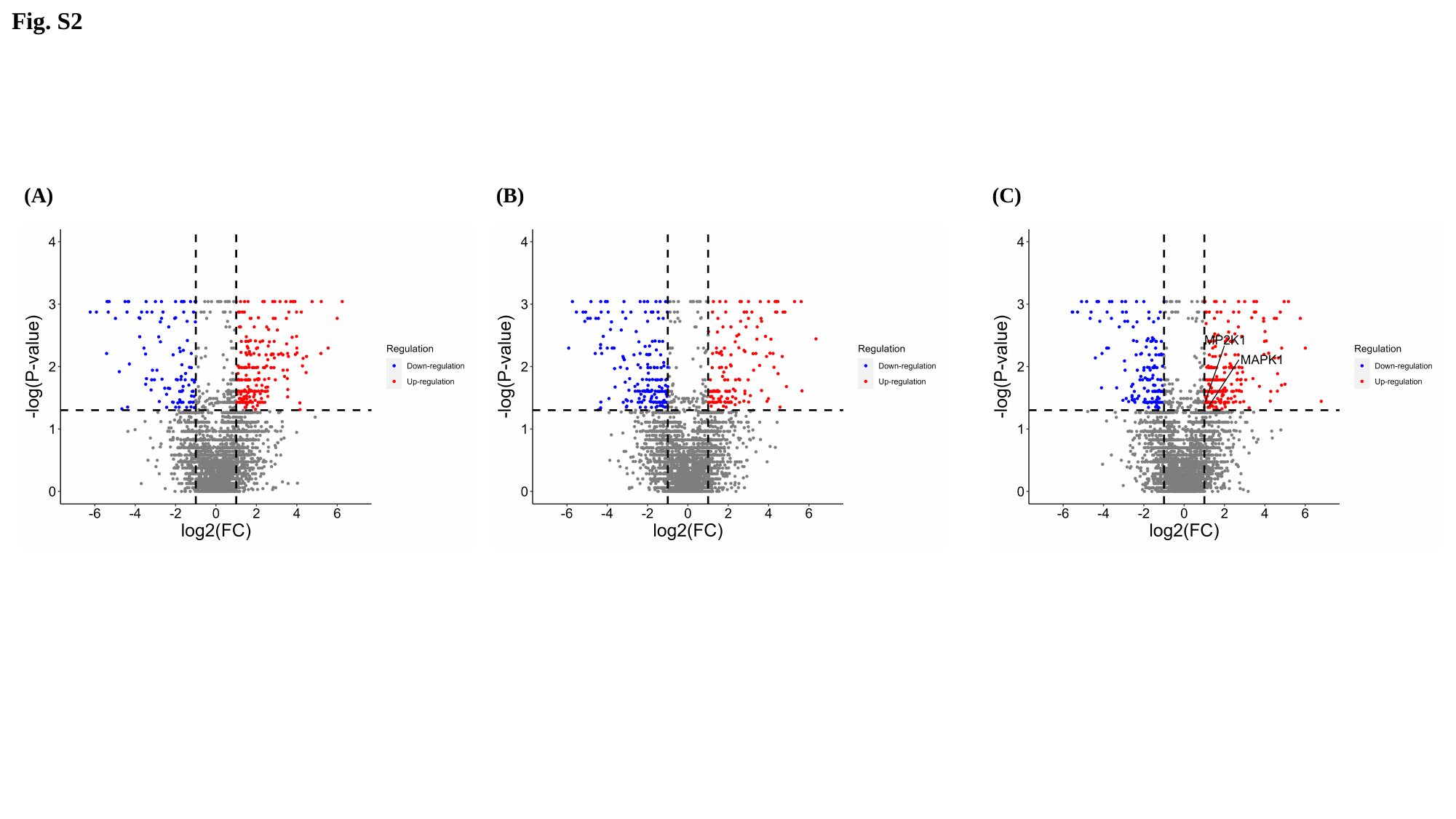

Fig. S2
(A)
(B)
(C)

### Slide 3
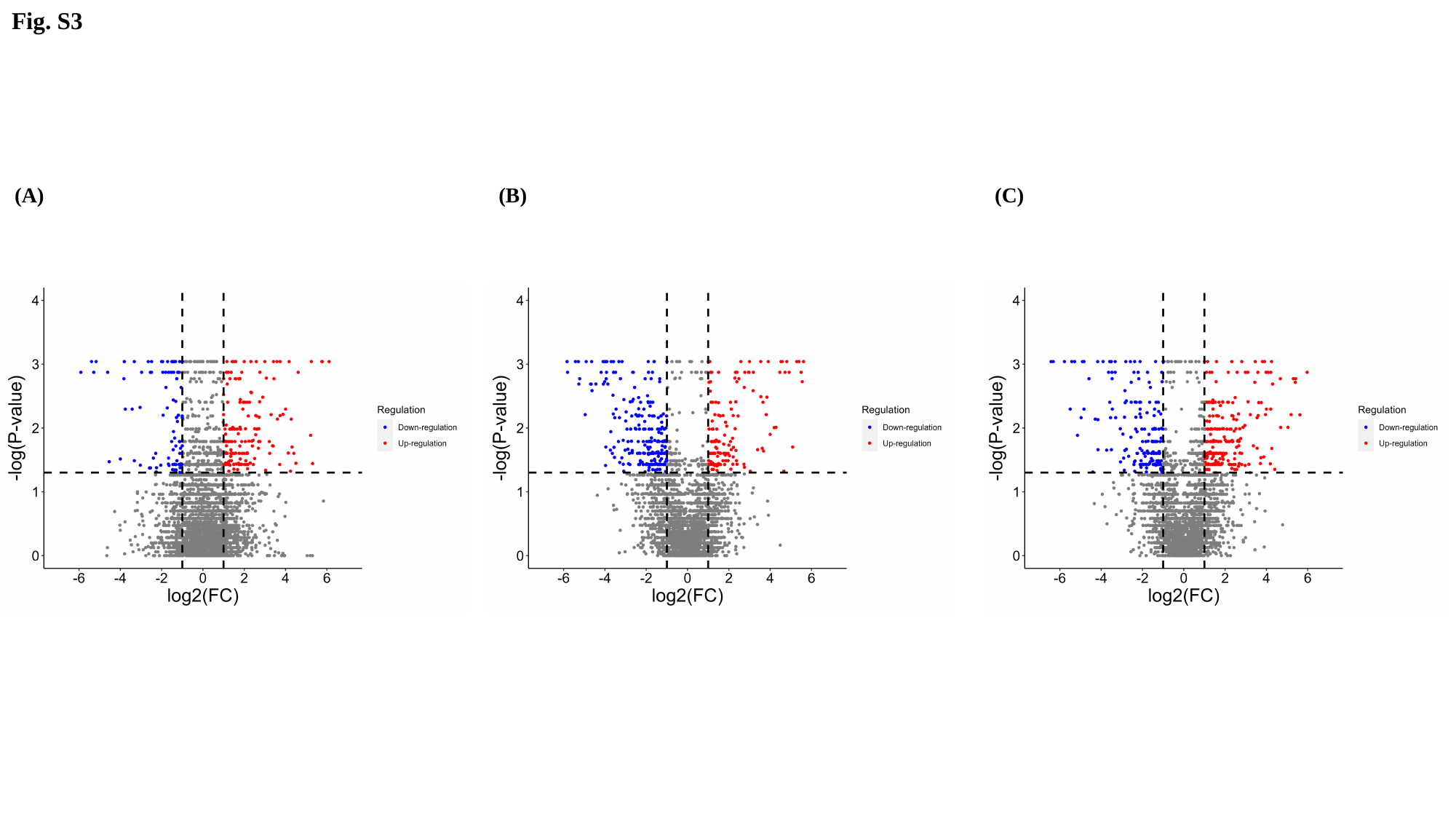

Fig. S3
(A)
(B)
(C)

### Slide 4
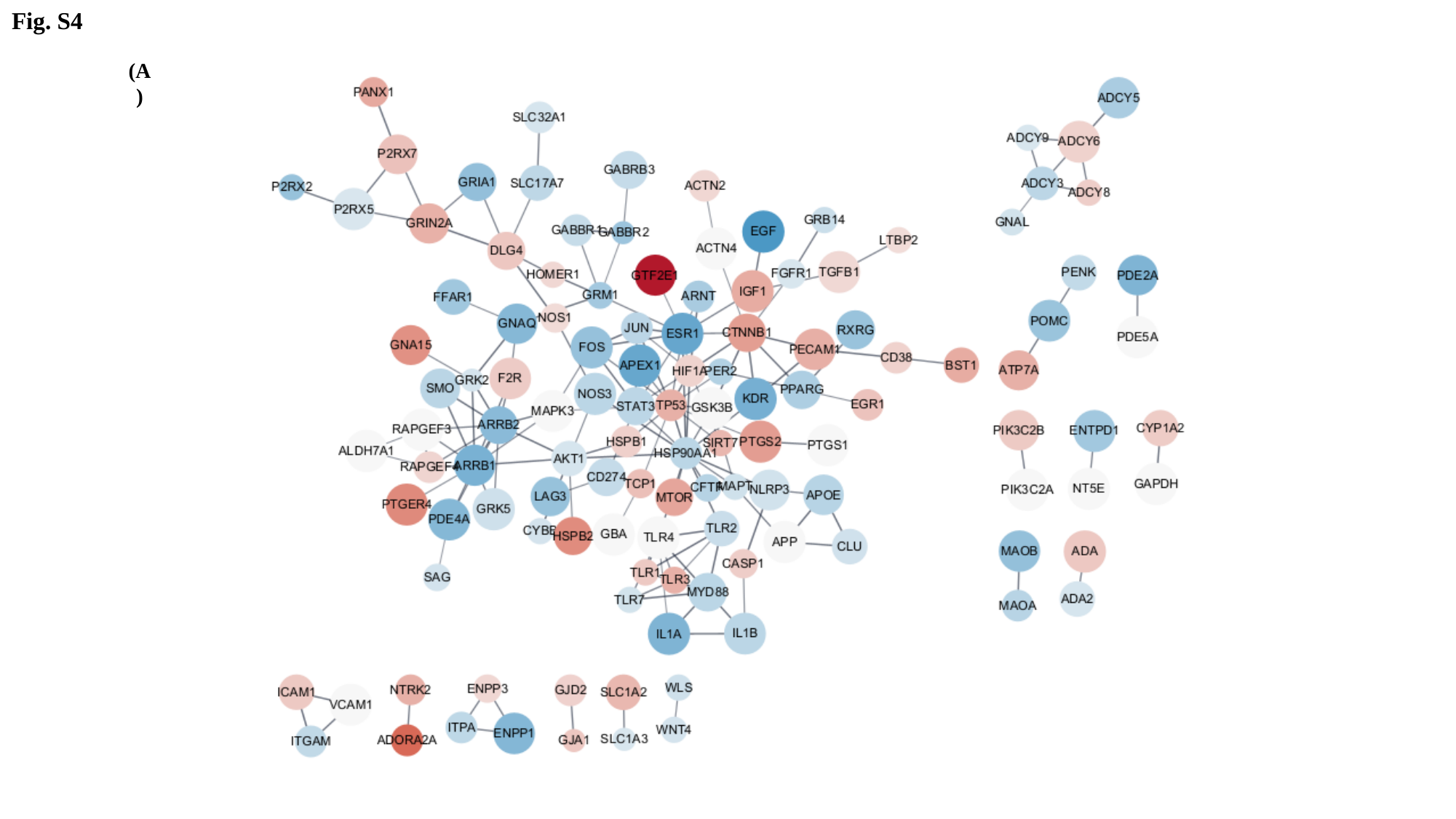

Fig. S4
(A)
(B)

### Slide 5
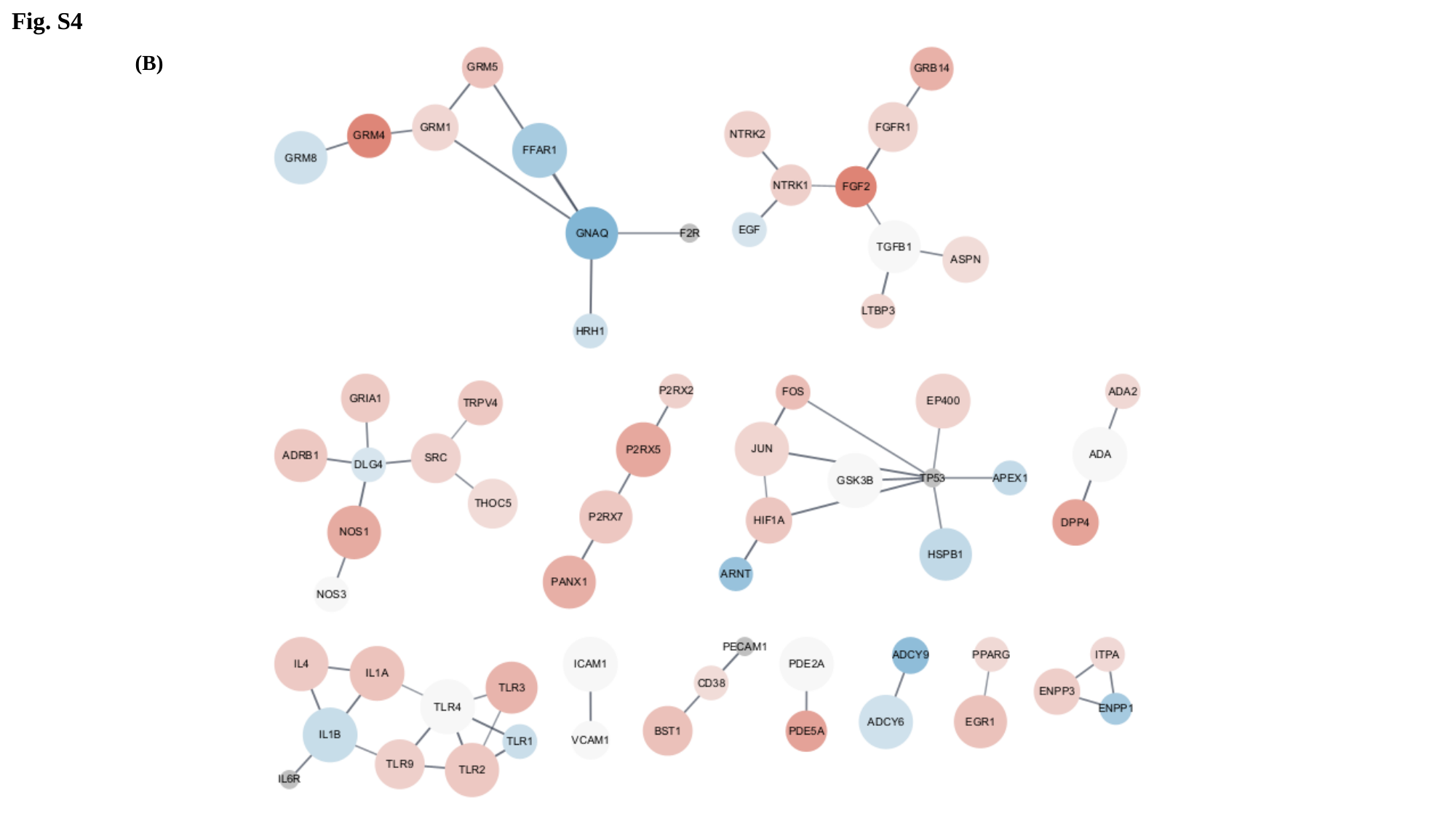

Fig. S4
(B)

### Slide 6
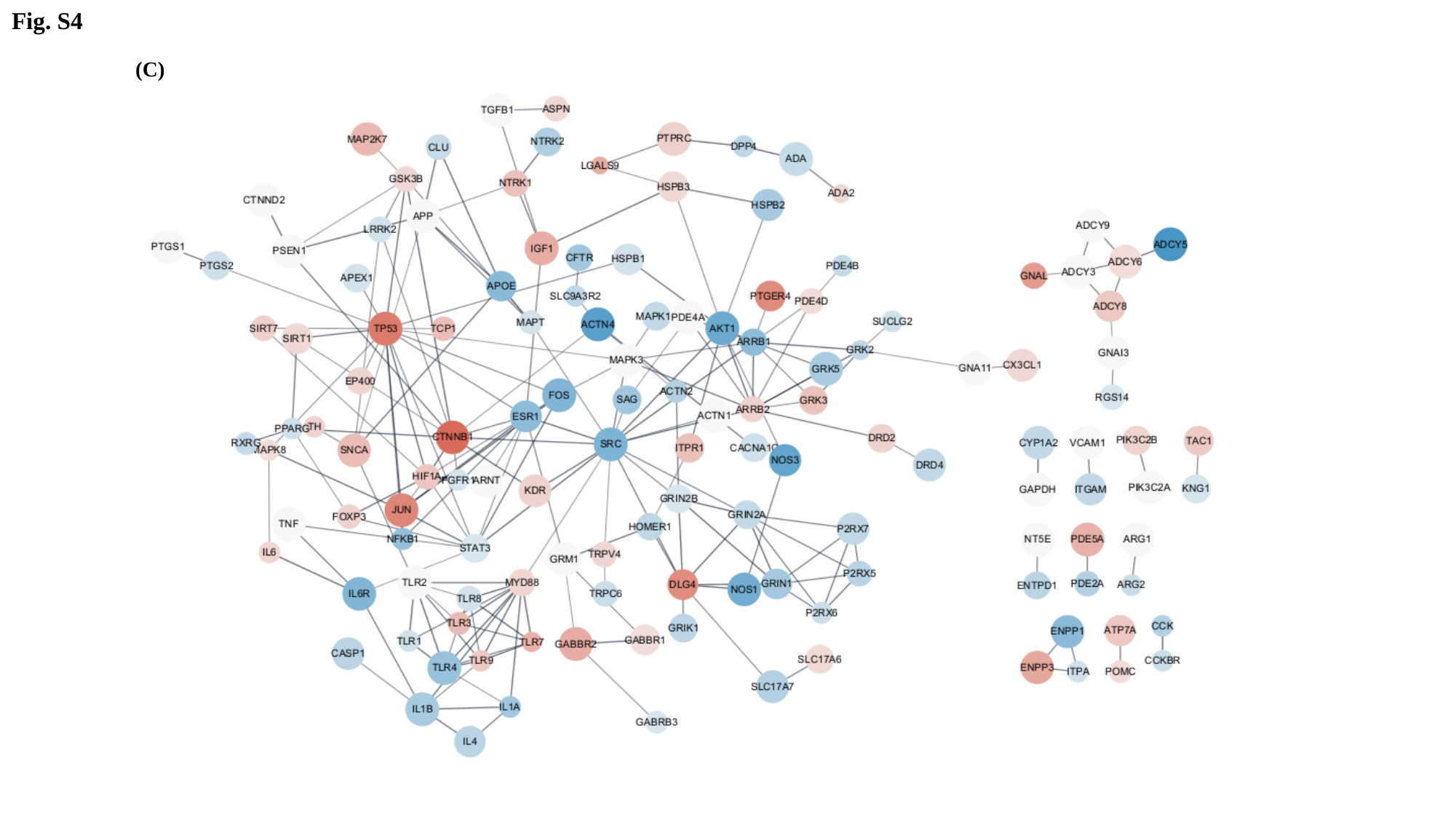

Fig. S4
(C)

### Slide 7
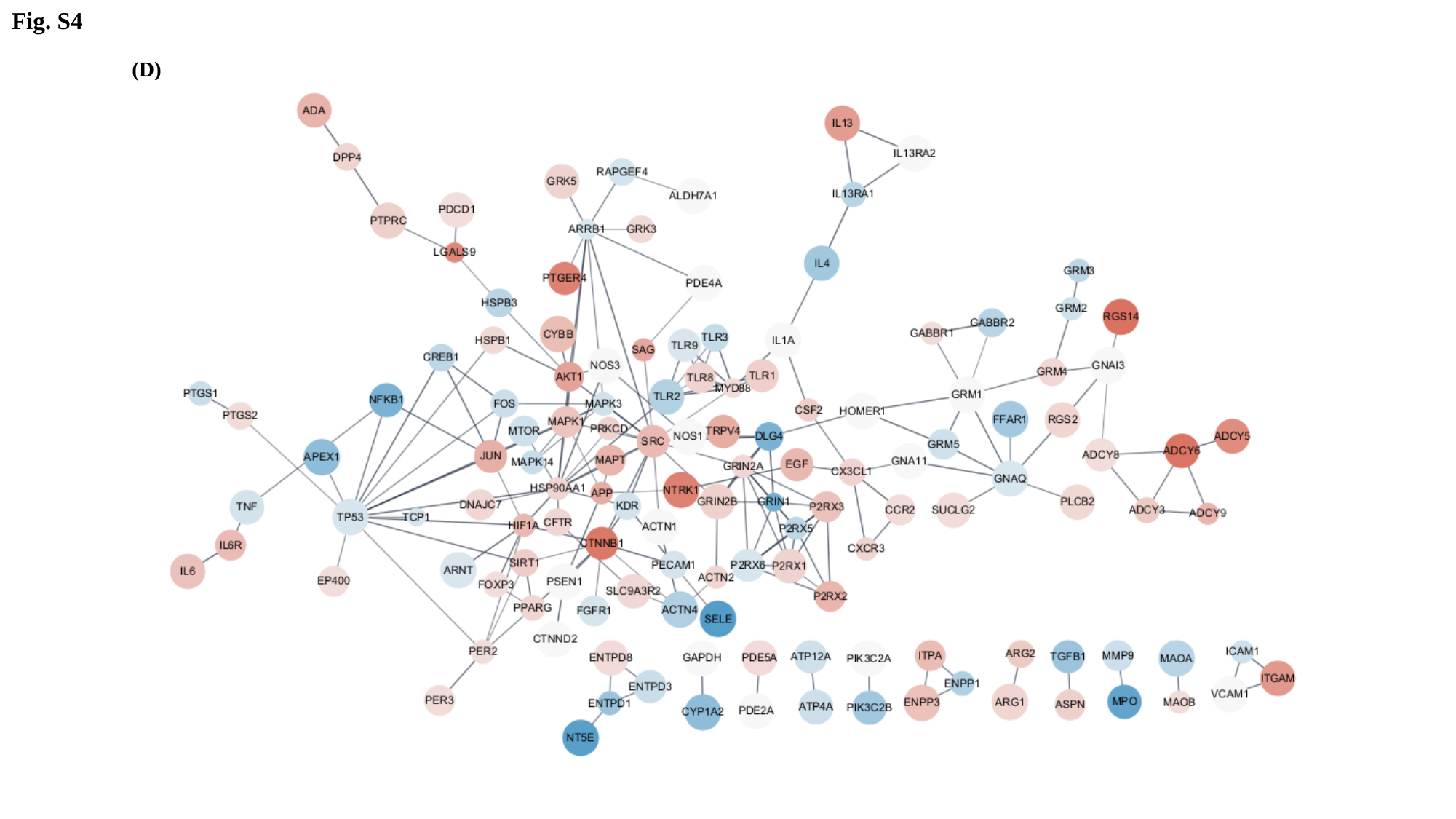

Fig. S4
(D)

### Slide 8
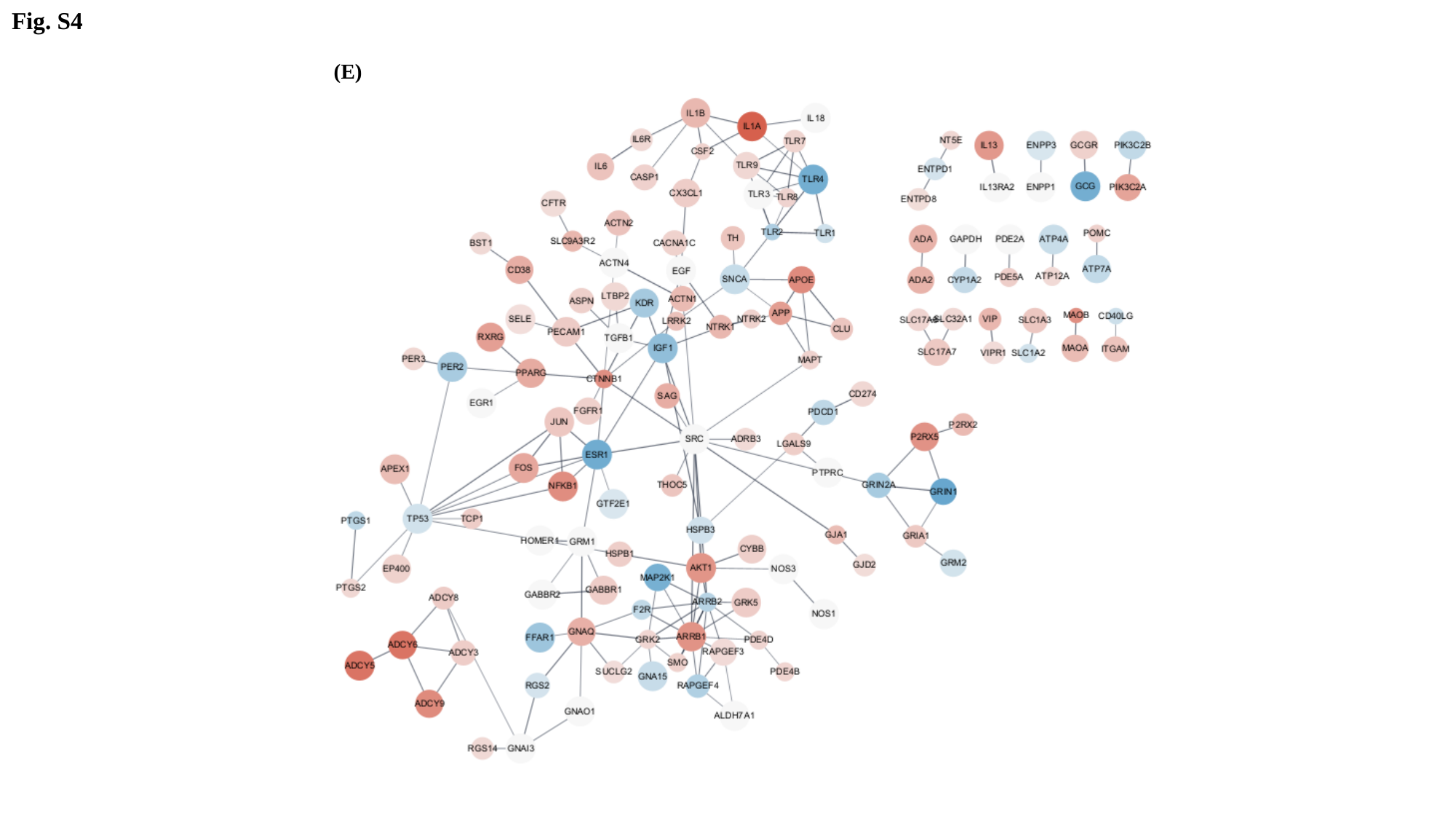

Fig. S4
(E)

### Slide 9
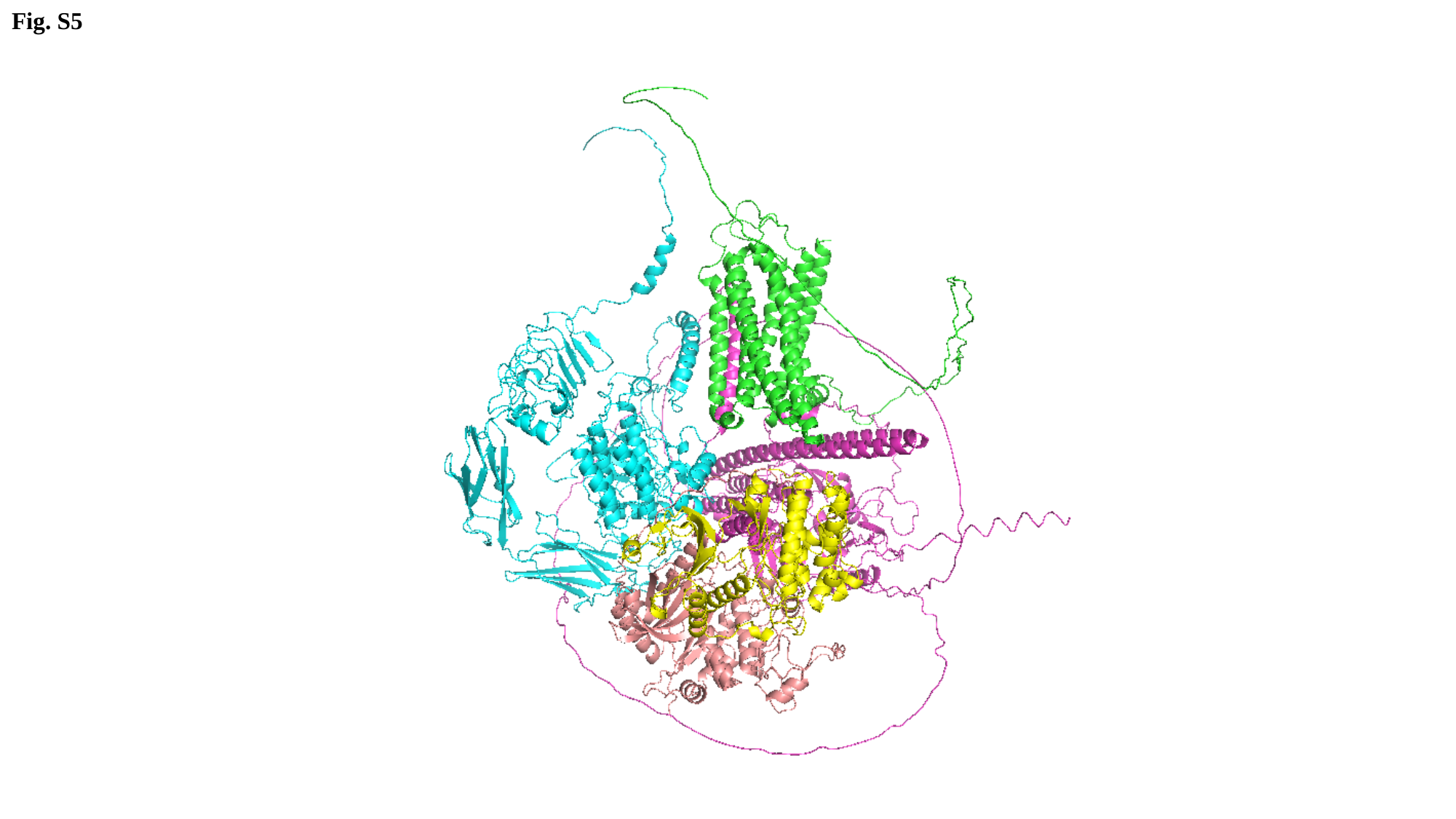

Fig. S5
