## Supplementary material for "Elucidating Interactome Dynamics of the A2A Adenosine Receptor": ADORA2A_supplementary_data: Supplementary_figure_legend.docx

**SUPPLEMENTARY FIGURES**

**Fig. S1. Workflow for Elucidating the Target of the ADORA2A Interactome via SRM-MS.**

The ADORA2A interactome was compiled from an extensive interactome databases, resulting in an initial selection of 513 proteins. Subsequently, this list was refined by selecting only peptides with a unique sequence, narrowing the selection to 502 proteins. Further refinement was achieved through SRM-MS analysis, culminating in the identification of 491 proteins and 3,852 peptides.

**Fig. S2. Differential Protein Expression of the ADORA2A Interactome in Comparison.**

Volcano plots illustrating the differential expression of normalized intensity in (A) ‘Salmon sperm vs. Flatfish sperm’, (B) ‘Flatfish sperm vs. Control group’, and (C) ‘Flatfish testis vs. Control group’. These plots highlight intensities exhibiting both a notable fold change (log_2_FC, x-axis) and high statistical significance (-log_10_ of *P*-value, y-axis). The dashed horizontal line represents the threshold for *P*-values (0.05), and the vertical line represents the threshold for fold change (2). The two vertical dashed lines demarcate the boundaries for downregulation and upregulation. Notably, MAPK1 and MAP2K1 are marked on the volcano plots to indicate their differential expression changes.

**Fig. S3. Comparative Volcano Plots of the ADORA2A Interactome.**

(A) Volcano plot illustrating the differential expression of transitions in the 'Salmon sperm vs. Control group' comparison. (B) Volcano plot depicting the differential expression of transitions in the 'Flatfish sperm vs. Flatfish testis' comparison. (C) Volcano plot presenting the differential expression of transitions in the ‘Salmon sperm vs. Flatfish testis’ comparison.

**Fig. S4. Network Analysis of Differentially Expressed Proteins Among Comparative Groups.**

(A) Network analysis depicting differentially expressed proteins in the ‘Flatfish sperm vs. Control group' comparison, showcasing 122 nodes and 9,168 edges. (B) Network analysis of differentially expressed proteins in the ‘Salmon sperm vs. Control group' comparison, featuring 64 nodes and 60 edges. (C) Network analysis illustrating differentially expressed proteins in the ‘Flatfish sperm vs. Flatfish testis' comparison, with 135 nodes and 215 edges. (D) Network analysis depicting differentially expressed proteins in the ‘Salmon sperm vs. Flatfish testis' comparison, consisting of 135 nodes and 200 edges. (E) Network analysis of differentially expressed proteins in the ‘Salmon sperm vs. Flatfish sperm' comparison, featuring 141 nodes and 179 edges. Red colors denote up-regulation, blue colors indicate down-regulation, and node sizes represent the *P*-value.

**Fig. S5. Predicted Interaction Sites within the Network Involving ADORA2A and Proteins Associated with Neuronal Differentiation in 5-Complex Model.**

(A) Predicted structure of proteins associated with neuronal differentiation depicted with Pymol. ADORA2A (green), NTRK1(cyan), APP (magenta), MAPK1 (yellow), and MAP2K1 (apricot) are present in the network.
